# Tracing information propagation in fish schools with a conditioned escape response

**DOI:** 10.64898/2026.08.04.742699

**Authors:** Vivek Jadhav, Cassandre Aimon, Fathimath Lamshana, Dari Trendafilov, Ramón Escobedo, Clément Sire, Guy Theraulaz, Vishwesha Guttal

**Affiliations:** Centre for Ecological Sciences, Indian Institute of Science, Bengaluru, Karnataka, 560012, India; Centre de Recherches sur la Cognition Animale, Centre de Biologie Intégrative, CNRS, Université de Toulouse – Paul Sabatier, Toulouse, France; Laboratoire de Physique Théorique, CNRS & Université de Toulouse – Paul Sabatier, France

## Abstract

Collective escape responses are widespread in animal groups, yet how threat-related information spreads from a few informed individuals to the collective remains poorly understood because the identities of the individuals who initially detect danger are rarely experimentally controlled. Here, we combined aversive conditioning, behavioural tracking, and computational modelling to investigate how escape information propagates through schools of *Puntigrus tetrazona*. We conditioned selected fish to associate a green light with an aversive stimulus. Subsequently, we tested the escape response of schools containing one informed and four naive fish to the green light. One informed fish was sufficient to trigger a collective escape response. Upon stimulus onset, conditioned fish accelerated and crossed the hurdle, after which naive fish sequentially increased their speed and followed. Temporal correlations in speed revealed a hierarchical leader–follower structure mostly aligned with hurdle-crossing order. Neither initial distance, viewing angle, nor relative orientation to informed fish predicted the order of the escape sequence. An agent-based model incorporating local alignment and distance regulation reproduced experimental observations and suggested that the informed individual likely remains partly attentive to its neighbours during escape. Overall, our work provides a general framework for experimentally dissecting information transfer in animal groups and identifying the behavioural mechanisms underlying collective escape dynamics.

## 1 Introduction

Group living and synchronised movement provide anti-predatory benefits [1]. Across taxa, predator detection by only a few individuals can trigger coordinated group-wide responses, generating propagating waves of motion that rapidly transmit information about danger through the collective [1, 2, 3]. These escape waves enable groups to respond cohesively even when most individuals never directly perceive the predator. In starling flocks attacked by falcons, escape waves originate near the point of attack and propagate away from the predator [4, 5]. In sulphur mollies, bird attacks trigger rapid diving manoeuvres that generate conspicuous surface waves [6]. Such coordinated escape responses can reduce predator success by confusing attackers [7, 8, 9], redirecting or disrupting predator attacks [6], or rapidly transmitting information about danger [10, 4].

A central question in this context is how information about a threat, available only to one or a few individuals, propagates socially within the group. Collective escape is part of a broader class of collective movement problems in which information held by one or a few individuals can influence the behaviour of an entire group. Previous work has shown that informed minorities can guide group movement toward resources or along migratory routes without explicit signalling or individual recognition [11, 12, 13, 14]. This is related to collective escape events where flocks or herds are pursued by a predator or chaser, where the external threat persists for a long duration, and the information may also be available to multiple individuals [15, 16, 17, 18]. However, threat-related information may also be localised and short-lived, requiring a transient behavioural response by informed individuals to be rapidly converted into social information [19, 20], as in the case of brief localised bird strikes on fish schools [6]. In such cases, inferring the precise quantitative mechanisms of information propagation remains difficult [21]. A key challenge is that during natural predator attacks, whether in the field or in controlled experiments, it is rarely possible to determine which individuals directly detect the predator and which respond only to social cues [22, 23]. This distinction, however, is crucial for separating direct responses to danger from socially mediated information transfer [24].

A powerful way to overcome this difficulty is to experimentally control which individuals initially respond to external perturbations [14, 25]. Here, we use an aversive conditioning protocol to trigger escape responses in selected individuals within a group [24]. In this setup, a green light, initially neutral, becomes an aversive conditioned stimulus after repeated pairing with a mild electric shock. By placing one conditioned fish among naive conspecifics, we can create a situation in which threat information is available only to a single individual. This experimental design allows us to track how the ‘private’ information held by an experimentally controlled initiator is transferred to the entire group through social interactions, a scenario that is difficult to quantify precisely in natural attacks.

We applied this approach to tiger barbs (*Puntigrus tetrazona*), a gregarious schooling fish. We conditioned individual fish to respond aversively to green light and then formed groups of five fish comprising one conditioned and four naive individuals. Using this system, we addressed three questions: (i) Can a single informed fish trigger a collective escape response? (ii) Which aspects of the informed fish’s escape behaviour are most important for initiating and propagating the escape response? (iii) What is the network of information transfer from the conditioned fish to the naive group members? To complement the experimental findings, we developed an empirically grounded computational model that reproduces the observed escape dynamics and provides a plausible mechanistic explanation for information propagation through the school.

## 2 Experimental setup and procedure

### 2.1 Study animals and conditioning procedure

The aversive conditioning protocol was adapted from [24] (for details, see electronic supplementary material, section S1). 14 fish were randomly selected and tagged with Visible Implant Elastomer at the base of the dorsal fin (Northwest Marine Technology, NMT Inc, Shaw Island, WA) following the manufacturer’s instructions and subsequently housed individually. These 14 fish were subjected to conditioning (hereafter referred to as conditioned individuals). Conditioning was conducted in a shuttle box (50 × 20 × 20 cm^3^) consisting of two compartments (20 × 20 × 20 cm^3^) separated by a 7.5 cm-high and 10 cm-wide trapezoidal hurdle positioned in the centre. Each compartment was equipped with 84 green light-emitting diodes (LEDs), which could be controlled independently (figure 1a,b). Two stainless steel mesh electrodes in each compartment, connected to a stimulator, delivered mild, controlled electric shocks. The current was set to the minimum intensity to elicit a visible behavioural response from the fish (7 V, 2.7 mA measured at the electrodes). Both visual and electrical stimuli were delivered manually. The box was filled with freshwater to a depth of 10 cm (2.5 cm above the hurdle).

**Figure 1:**
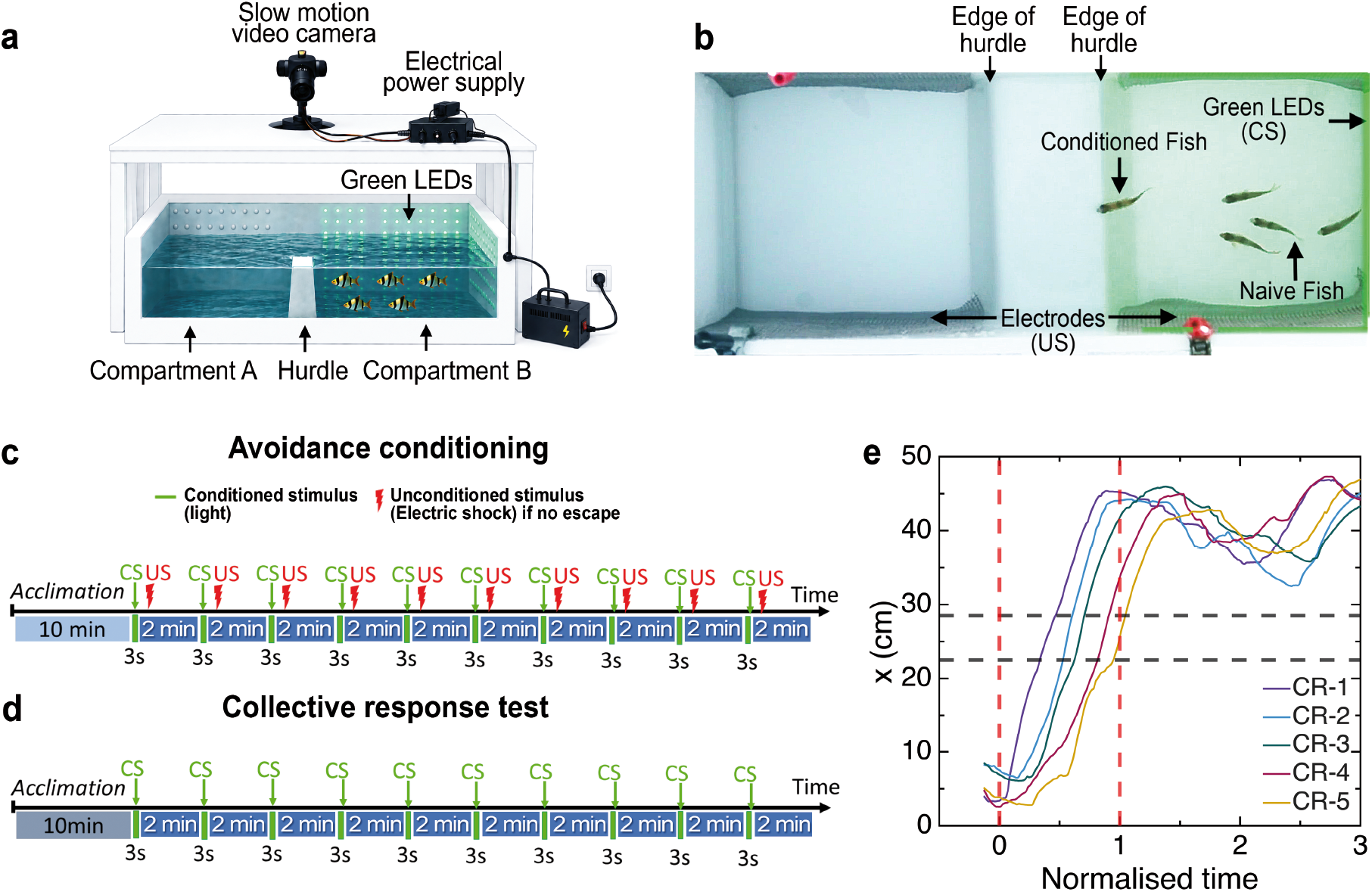
Experimental setup and protocol. **(a,b)** Shuttle box used for aversive conditioning and avoidance testing. **(c)** Conditioning procedure and **(d)** experimental protocol. **(e)** Trajectories of individual fish from a representative trial. The vertical red dotted lines show the limits of the escape phase, where *t* = 0 corresponds to the onset of the green light and *t* = 1 to the time at which the last fish completely crossed the hurdle. The horizontal black dashed lines represent the trapezoidal hurdle. The hurdle position is slightly offset (22.5 cm and 28.5 cm instead of 20 cm and 40 cm) to account for tracking errors and to facilitate identification of crossing order.

Each conditioning session consisted of 10 trials separated by a 2 min inter-trial interval. During each trial, the conditioned stimulus (CS; green light) was presented for 3 sec in the compartment occupied by the focal fish. If the fish failed to cross the hurdle within this period, the unconditioned stimulus (US; electric shock) was delivered. A trial was considered successful if the fish crossed the hurdle before the onset of the US (i.e., within 3 sec of the onset of the CS; figure 1c). Fish underwent repeated conditioning until reaching a performance criterion of 80% (i.e., at least 8 successful trials out of 10 without receiving the US). Only individuals meeting this criterion were used in subsequent collective escape experiments (7 out of 14 fish).

### 2.2 Experimental protocol

Test groups were composed of one conditioned and four naive fish. Each trial consisted of a 3-second presentation of the conditioned stimulus, during which collective behaviour was recorded. Trials were separated by a 2-minute inter-trial interval (figure 1d). The order in which the conditioned fish were tested with fresh naive fish was randomised.

As a control experiment, 6 groups of 5 naive fish were tested in the shuttle box without electric shocks to quantify the baseline escape response induced by the green light alone.

Finally, fish trajectories were extracted using a custom tracking software specifically developed at CRCA.

### 2.3 Quantification of collective escape response

We partitioned each trial into three phases, *Initial, Escape*, and *Relaxation*, to compare the group properties and interaction structure before, during, and after the escape response:

- *Initial phase*: From the start of the trial to the onset of the green light. This phase provides a baseline for schooling behaviour under unperturbed conditions. During this phase, no systematic differences are expected between conditioned and naive fish in terms of swimming speed (electronic supplementary material, figure S1).
- *Escape phase*: From the onset of the green light to the time at which the last fish has completely crossed the hurdle (see figure 1b). This phase captures the collective escape response.
- *Relaxation phase*: From the end of the escape phase to the end of the trial. This phase corresponds to the transient return to baseline behaviour. As in the initial phase, no systematic differences are expected between the conditioned and naive fish in terms of speed (electronic supplementary material, figure S1).

We normalise time such that *t <* 0 corresponds to the initial phase, 0 ≤ *t* ≤ 1 corresponds to the escape phase, and *t >* 1 corresponds to the relaxation phase. This normalisation enables direct comparison of escape dynamics across trials. We define the crossing rank (CR) as the order in which individuals first cross the edge of the hurdle encountered along their escape trajectory, with the first fish to cross (the conditioned fish) assigned rank 1, and the last fish assigned rank 5.

We quantified collective behaviour using two standard metrics [26] (for details, see electronic supplementary material, section S1):

- *Group dispersion* (*D*) quantifies spatial cohesion of the group and is defined as the mean distance of individuals from the group barycentre. Similarly, *D*_*x*_ and *D*_*y*_ denote the mean distances from the barycentre along *x*- and *y*-axes, respectively. We also computed dispersion without the conditioned fish (*D*_wc_).
- *Group polarisation* (*P*) quantifies the degree of alignment among fish, with *P*_*x*_ and *P*_*y*_ representing alignment along *x*- and *y*-axes.

## 3 Results

### 3.1 Conditioned fish elicits collective escape responses

In control groups, no fish crossed the hurdle in 58 out of 63 (92%) trials, indicating that the green light alone does not reliably elicit crossing behaviour (electronic supplementary material, figure S2). In contrast, for groups with one conditioned fish, all five fish crossed in 87 of 125 trials, with only 4 trials eliciting no response from any fish (electronic supplementary material, figure S2). Of these 87 trials in which all five fish crossed the hurdle, we removed 34 trials from our analysis due to technical issues during the trials, such as difficulty tracking the fish, background disturbances in the experimental room, or aggressive interactions among a few fish before the stimulus was presented. Thus, we have 53 trials to study the collective escape response, in which the conditioned fish initiated the response by crossing the hurdle first, followed by all naive individuals.

Figures 2a,b,c show representative time series of individual speeds, group polarisation, and group dispersion, respectively, for the trial illustrated in figure 1e. During the initial phase, individuals swim at an average speed of approximately 5 cm/s (≈ 1.3 BL/s), and the school is cohesive and remains highly polarised. Following the onset of green light, the conditioned fish (CR = 1) rapidly increases its speed and crosses into the adjacent compartment. The naive individuals subsequently accelerate and cross the hurdle in sequence, following the conditioned fish. This sequential increase in speed mirrors the order in which the individuals cross. During the escape phase, group polarisation transiently increases, whereas group cohesion decreases, resulting in higher dispersion. Once all fish have crossed the hurdle and reached the adjacent compartment, swimming speed, polarisation, and dispersion progressively return to values comparable to those observed in the initial phase.

**Figure 2:**
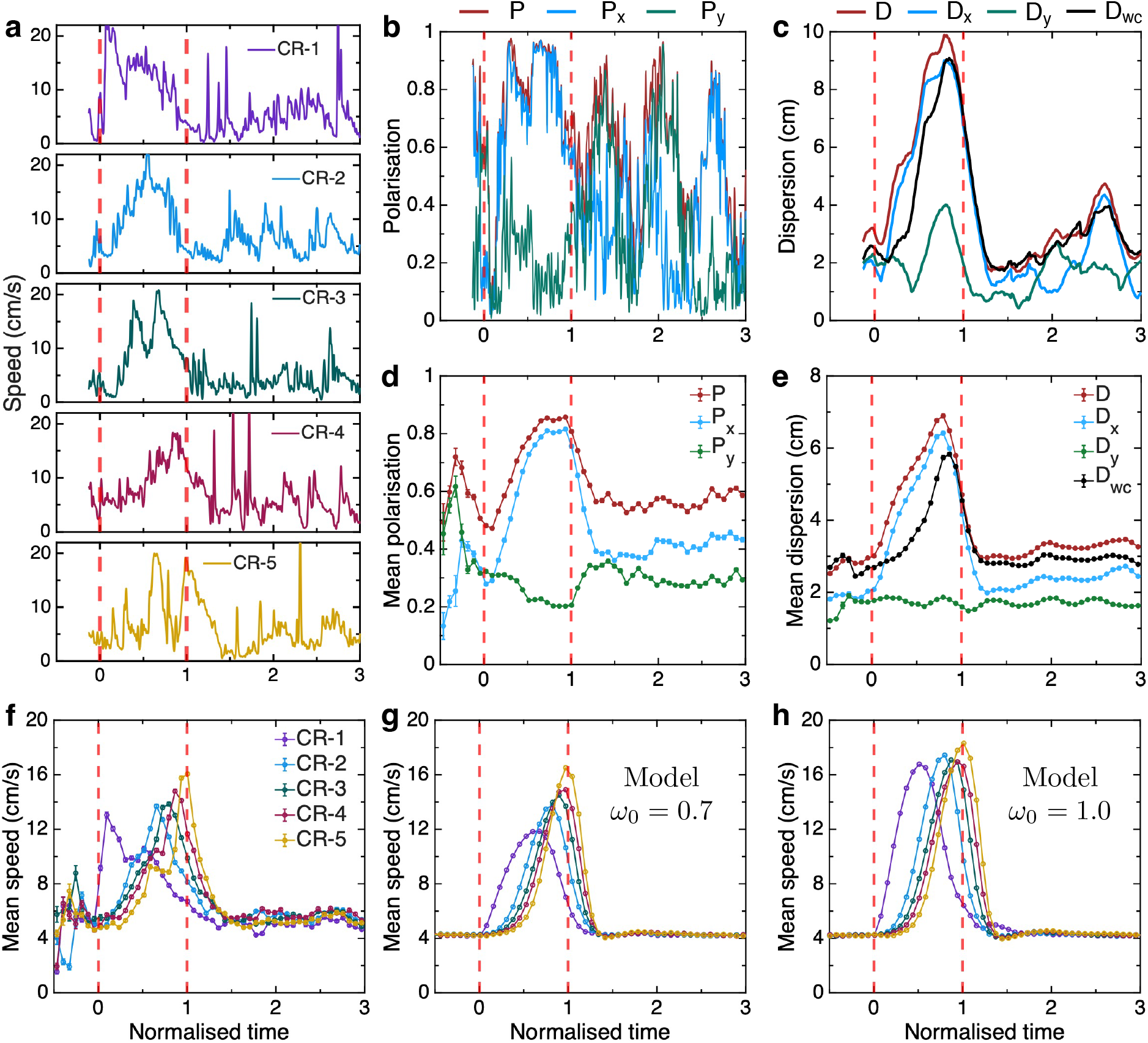
Dynamics of speed, polarisation, and dispersion during collective escape. Time series of **(a)** individual speed of fish sorted by crossing rank **(b)** polarisation *P* (*t*), **(c)** dispersions *D*(*t*) and *D*_wc_(*t*), (from trial shown in figure 1e), **(d)** mean polarisation and **(e)** mean dispersion across trials, and **(f)** mean individual speed sorted by crossing rank. **(gh)** Influence of escape-force strength on mean speed during collective escape dynamics in numerical simulations, for *ω*_0_ = 0.7 and *ω*_0_ = 1.0. Error bars indicate standard errors, red dashed lines indicate green light onset and crossing time of last fish.

To generalise across experiments, we computed the average polarisation, dispersion, and speed as functions of normalised time *t* (figure 2d,e,f). Both *P* (*t*) and *D*(*t*) increase during the escape phase (*t* ∈ [0, 1]). As fish move from one compartment to the other along the *x*-axis, both *P*_*x*_ and *D*_*x*_ increase during the escape phase, thereby contributing to the overall increase in group polarisation and dispersion, respectively. In contrast, *P*_*y*_ and *D*_*y*_ remain largely unchanged throughout the escape phase. We further show that the increased polarisation may arise, at least partly, from the geometry of the experimental tank, as it constrains fish movement primarily along a single spatial axis (electronic supplementary material, section S3 and figure S3a,b). Moreover, the dispersion computed after excluding the conditioned fish *D*_wc_ also increases during the escape phase, showing that naive fish cross the hurdle sequentially rather than as a subgroup.

We find that fish accelerate sequentially in the order they cross the hurdle. Moreover, the average maximum speed of fish v^max^ increases with crossing rank *i*, i.e., 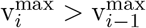 (although 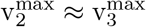, figure 2f). Thus, as more fish cross into the adjacent compartment, the remaining fish tend to swim faster to rejoin the group. Once all fish have crossed the hurdle, polarisation, dispersion, and swimming speed return to values comparable to those observed during the initial phase. Although no persistent differences in schooling behaviour are expected between the initial and relaxation phases, a transient adjustment period may occur immediately after the escape response before the school settles back into a stable collective state. To assess this possibility, we computed the probability density functions of polarisation, dispersion, and individual speed across each of the three phases. Group-level polarisation, dispersion, and swimming speed were indeed similar in the initial and relaxation phases (figure 3). We note, however, that the initial phase contains fewer data points than the relaxation phase, resulting in noisier probability density estimates.

**Figure 3:**
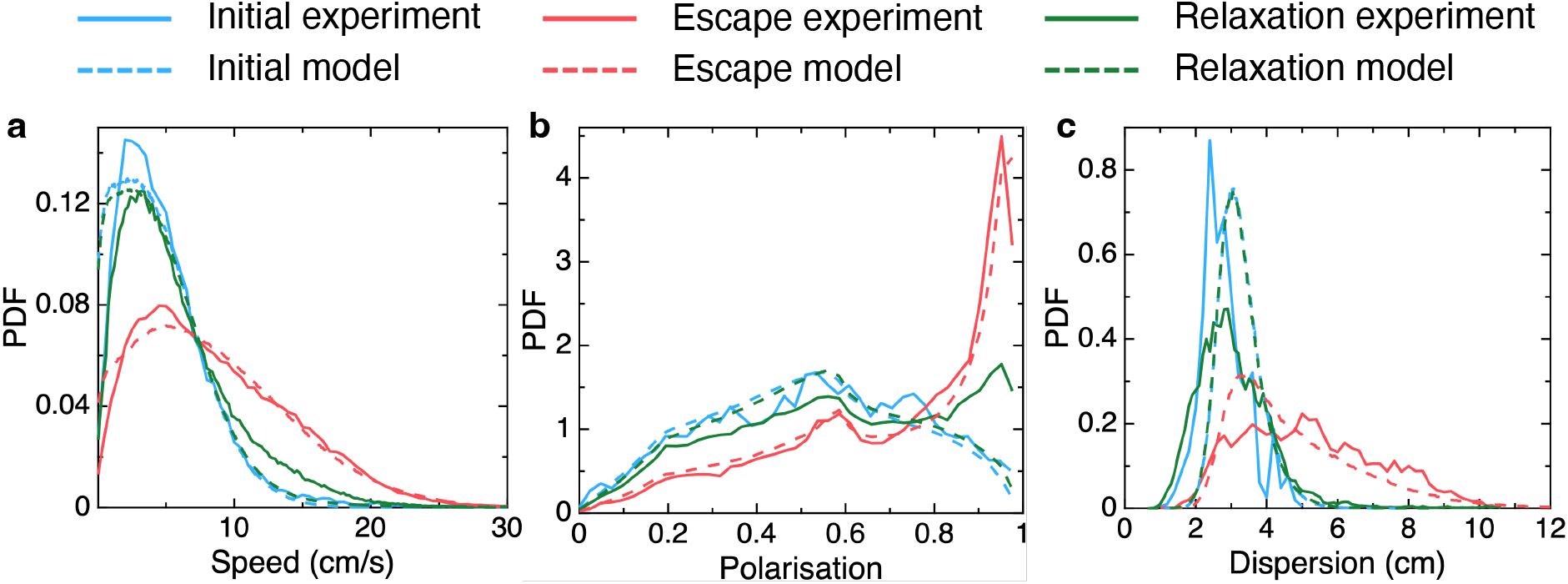
Probability density functions (PDF) of speed, polarisation, and dispersion across the three phases. **(a)** Individual swimming speed. **(b)** Polarisation. **(c)** Dispersion. Blue, red, and green lines correspond to the initial, escape, and relaxation phases, respectively. Mean ± SD values for experimental data are: ⟨v⟩_initial_ = 5.25 ± 4.25 cm/s, ⟨v⟩_escape_ = 9.02 ± 5.86 cm/s, ⟨v⟩_relaxation_ = 6 ± 4.5 cm/s; ⟨*P* ⟩_initial_ = 0.53 ± 0.25, ⟨*P* ⟩_escape_ = 0.70 ± 0.25, ⟨*P* ⟩_relaxation_ = 0.58 ± 0.25; and ⟨*D*⟩_initial_ = 3.04 ± 0.68 cm, ⟨*D*⟩_escape_ = 5.4 ± 1.9 cm, ⟨*D*⟩_relaxation_ = 3.12 ± 1.02 cm.

To investigate whether crossing order was related to the initial spatial configuration of the group, we quantified the distance between each naive fish and the conditioned individual at *t* = 0. We found no significant relationship between crossing rank and the initial distance to the conditioned fish (electronic supplementary material, section S4 and figure S4a). In addition, we measured the viewing angle *ψ*_*i*1_, defined as the angular deviation required for fish *i* to orient toward the conditioned fish, and the relative orientation *ϕ*_*i*1_, defined as the difference in heading angle between fish *i* and the conditioned one, for *i* = 2, …, 5. Neither *ψ*_*i*1_ nor *ϕ*_*i*1_ had a significant effect on the crossing rank (electronic supplementary material, section S4 and figure S4b,c). These results indicate that the initial spatial position and orientation of naive individuals relative to the conditioned fish do not predict the order in which they cross the hurdle during the collective escape.

We then quantified the crossing time of each individual 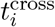, defined as the latency between the onset of the green light and the instant at which fish *i* crosses the edge of the hurdle along its escape trajectory, where *i* denotes the crossing rank. Crossing latency increased linearly with crossing rank, with the conditioned fish and the last fish crossing the hurdle, on average, 0.39 and 0.9 normalised time units after stimulus onset, respectively (figure 4a, see electronic supplementary material, figure S5a for time in seconds). The first naive fish to follow the conditioned fish crossed after an average delay of 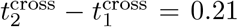 normalised time units. This inter-crossing interval decreased to ≈ 0.11 normalised time units between the first and second naive fish, and remained constant for subsequent crossings (figure 4b, see electronic supplementary material, figure S5b for time in seconds). These results show that the delay between successive crossings does not decrease as more fish cross the hurdle.

**Figure 4:**
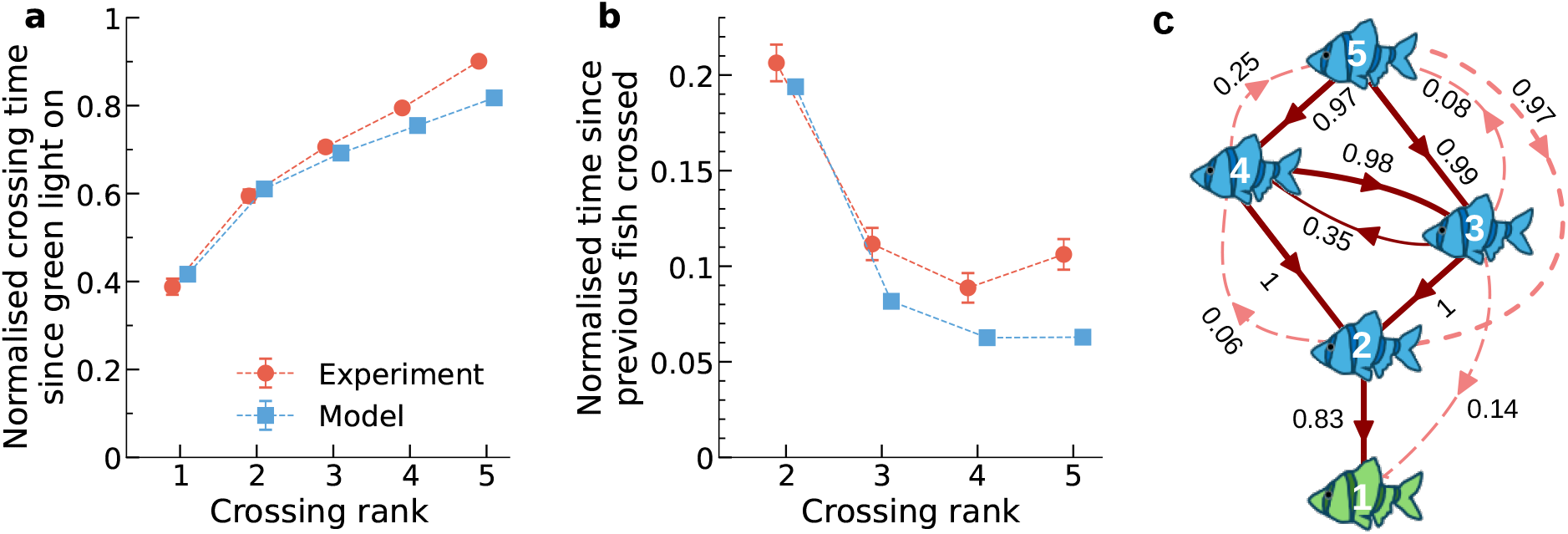
Sequential escape response and leader–follower interaction networks during collective escape. (**a**) Mean normalised latency to cross the hurdle for each crossing rank measured from the onset of the green light stimulus. (**b**) Mean normalised time between successive crossings. The first (2nd, 3rd,…) naive fish crossing the hurdle follows the conditioned fish and has rank 2 (resp. 3, 4,…). Blue symbols correspond to experimental data, red symbols to model simulations. Error bars indicate the standard error. Symbols are shifted horizontally to avoid overlap. (**c**) leader–follower network: Nodes correspond to fish identified by crossing rank (white number). Directed edges indicate leader–follower relationships and point from followers to leaders. Dark arrows correspond to leader–follower pairs reconstructed from experimental data, all of which are reproduced by the model simulations. Edge labels indicate the fraction of simulations in which the pairwise correlation between those two nodes was statistically significant. Additional dotted lines correspond to leader–follower relationships observed only in model simulations, not in the experimental data.

### 3.2 Leader–follower relationships and information propagation

As described above, conditioned fish rapidly crossed the hurdle upon the onset of the green light, with naive fish following soon after, suggesting that the conditioned fish initiates the collective response through a rapid change in swimming speed. The subsequent propagation of this behavioural perturbation is then governed by the local interaction rules between neighbouring fish. Although the information may be propagated through changes in orientation [3, 27, 28, 29, 4], speed [3, 30] or both, the geometry of our experimental setup constrains movement primarily along a single spatial axis, thereby confounding directional information transfer with the effects of spatial confinement. Therefore, we focus on speed-mediated information transfer rather than orientation-mediated information transfer.

To quantify leader–follower relationships during escape, we compute the cross-correlation of individual swimming speeds, which measures the degree to which the speed fluctuations of a fish influences that of another fish after a time delay *τ* . Given the speed time series v_*i*_(*t*) and v_*j*_(*t*) of fish *i* and *j*, respectively, the normalised cross-correlation at lag *τ* is defined as:

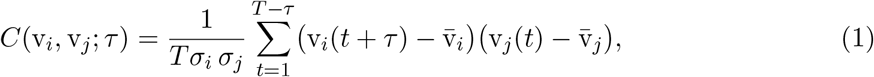

Where 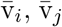, and *σ*_*i*_, *σ*_*j*_ are the mean and standard deviation of each speed time-series computed over the time window (*t* = 1, …, *T*) of an escape phase. We then define the lag *τ*_*ij*_ as the value of *τ* at which *C*(v_*i*_, v_*j*_; *τ*) is maximum. Individual *i* is considered to lead *j* when *τ*_*ij*_ *<* 0, and vice versa. The values *τ*_*ij*_ for all pairs (*i, j*) allow us to construct a leader–follower network for each trial [18, 31, 12, 32]

The interpretation of leader–follower relationships based on the temporal correlation function defined in equation 1 applies only to stationary time series. However, the speed time series recorded during the escape phase are clearly non-stationary, being characterised by rapid accelerations followed by decelerations. When this method is applied to speed trajectories from the null model, where all individuals respond independently to an external stimulus without interacting with each other, it produces a spurious hierarchical network that closely mimics the data [33] (see electronic supplementary material, section S3, figure S6, and figure S7). Therefore, to correctly infer leader–follower relationships, we employ a permutation test, a nonparametric test, to reliably assess the significance of correlations in non-stationary time series [34, 35, 36] (see electronic supplementary material, section S5 for details). The principle underlying this method is that to establish speed correlations between two fish, correlations computed between two fish from the same trial should be greater than those obtained between fish from different trials. We note that this method correctly infers the absence of interaction between individuals in the null model; see electronic supplementary material, figure S8.

We display the resulting leadership network in figure 4c, in which each node represents a fish characterised by its crossing rank, and directed edges from followers to leaders are drawn as solid lines. First, we expected no behavioural hierarchy during the initial and relaxation phases. Indeed, as expected, we find no leader–follower pairs during these phases. Next, we expected that the conditioned fish might influence all fish during the collective escape phase, owing to the small group size and the constrained geometry of our experiments. In contrast, we observe that the conditioned fish significantly influences only a single naive fish, which is also the first to cross the barrier. Overall, the network reveals a hierarchical leadership structure closely aligned with crossing order (also see electronic supplementary material, table S1). In particular, the fish with crossing rank 2 typically followed the conditioned fish, and the fish with crossing rank 3 followed the fish with rank 2. Consequently, edges are oriented from higher to lower crossing ranks, indicating a sequential propagation of information through the group. Exceptions involve fish with crossing ranks 3 and 4, for which significant correlations were detected in both directions.

### 3.3 Modelling collective escape dynamics

We developed an agent-based model to investigate the behavioural mechanisms underlying the propagation of threat information from conditioned fish to naive fish. Our model builds on the general framework of self-propelled collective motion models and adapts the framework proposed in [37].

#### Model rules

The model describes the motion of conditioned and naive fish through coupled equations for speed and heading. The complete mathematical formulation, interaction rules, simulation protocol, and parameter values are provided in the electronic supplementary material, section S2. For completeness, we briefly describe the model here. Each fish experiences four forces: self-propulsion, speed matching and alignment with neighbours, distance regulation, and wall avoidance. When the green light is turned on, only the conditioned fish experiences an escape force (directed toward the centre of the adjacent compartment), whose relative strength is controlled by *ω*_*t*_ ∈ [0, 1]. Here, *ω*_*t*_ = 0 for *t <* 0, and takes a constant value *ω*_0_ from the time the light is turned on to the time the conditioned fish crosses the hurdle, and decays exponentially thereafter. Note that *ω*_0_ = 0 corresponds to the absence of escape force, whereas *ω*_0_ = 1 corresponds to the absence of social interactions with neighbours.

Simulations were run using boundary conditions identical to the experimental tank. We varied the strength of the escape response (*ω*_0_) to investigate its effect on collective escape dynamics. Furthermore, to evaluate the roles of social interactions and geometric constraints, we considered a null model in which individuals experience no social force. In this null model, all fish experience an escape force once the light is turned on (see electronic supplementary material, section S3).

#### Model results

With the chosen parameter values, the model simulations qualitatively reproduce the main features observed experimentally (electronic supplementary material, figure S9). PDFs of polarisation, dispersion, and swimming speed are in agreement with the empirical ones (figure 3). The null model shows that the increase in polarisation during the escape phase arises primarily from the geometric constraints imposed by the experimental tank, rather than from collective coordination (electronic supplementary material, figure S3a). While the dispersion in the escape phase is qualitatively similar in the null model, the group is less cohesive during the initial and relaxation phases due to the absence of social interactions among individuals (electronic supplementary material, figure S3b). However, the geometric constraints of the experimental tank did not significantly influence any other measured group property (electronic supplementary material, figure S3).

Figure 4 shows that the sequential propagation of the escape response is also qualitatively well reproduced: the empirical latencies increase linearly with the crossing rank (figure 4a), the inter-crossing interval decreases for the first naive fish and remains approximately constant for the following fish (figure 4b), and the leader–follower network is predominantly oriented from higher to lower ranks, as observed experimentally (figure 4c). Only occasionally weak reverse connections appear (e.g., edges from rank 4 to rank 5, in 25% of simulations), but consistent with the bidirectional links observed between crossing ranks 3 and 4 in the experimental data. The model also confirms that crossing order is independent of the initial spatial configuration relative to the conditioned fish: neither the distance to the conditioned fish, the viewing angle, nor the relative orientation, predicts which naive fish will cross the hurdle after the conditioned fish (electronic supplementary material, figure S4).

We then investigated the effect of the initial value of the escape-force weight *ω*_0_ on the mean speed profiles during crossing events. Our aim was twofold: first, to identify the value of *ω*_0_ that best reproduces the empirical speed profiles shown in figure 2f, and second, to understand how the escape force and social interactions balance during the escape response.

We found that simulations with *ω*_0_ = 0.7 best reproduced the shape of the mean speed profiles, both in the amplitudes and the locations of the peaks, and also in the order across crossing ranks (figure 2g). When *ω*_0_ = 1, the mean speeds are higher than when *ω*_0_ = 0.7 (figure 2h). The higher speeds observed at larger values of *ω*_0_ result from the combination of two mechanisms. When *ω*_0_ increases, the net force acting on the conditioned fish also increases because the gain in the escape force outweighs the loss in the social force, and this results in a higher speed. In turn, when *ω*_0_ is smaller, social interactions with the naive ones are more intense and contribute more to slow down the escape of the conditioned fish (electronic supplementary material, figure S10).

Examining the effect of *ω*_0_ on the structure of the leader–follower network, we found that, when *ω*_0_ adopts values for which the conditioned agent retained a significant degree of social coupling with the naive ones (*ω*_0_ = 0.7), the fraction of simulations in which the fish with crossing rank 2 significantly followed the conditioned fish is 1.5 times higher than when this coupling is reduced: from above 0.8 when *ω*_0_ = 0.7 to about 0.5 when *ω*_0_ = 0.8, remaining at similarly low values, 0.2 and 0.1 for *ω*_0_ = 0.9 and *ω*_0_ = 1 respectively (electronic supplementary material, figure S11). This sharp drop suggests that, although the social coupling 1 − *ω*_*t*_ can be initially small relative to the escape force, a minimal degree of social coupling between the conditioned and the naive agents must be maintained during escape to reproducing the experimentally observed propagation dynamics.

## 4 Discussion

Understanding how information about threats propagates through animal groups requires correctly identifying the individuals who detect the threat and studying how this private information, often initially available only to a few individuals, is converted into group-level social information, thereby leading to a collective escape [21]. Here, we addressed this question by precisely controlling which individuals respond to a short-lived aversive stimulus and by analysing how this information subsequently spreads through the group. First, we showed that in a school of five fish, a single conditioned individual is sufficient to trigger a collective escape response [11]. Following the onset of green light, conditioned fish rapidly increased their swimming speed and crossed the hurdle into the adjacent compartment. Naive individuals subsequently increased their speed and followed, indicating that threat-related information is likely transmitted through local changes in swimming speed. Our key result is that briefly available threat information detected by the informed individual can lead to a sequential, socially mediated escape response at the group level. Thus, in our experiments, collective escape emerges from individual escape driven by the private information of the conditioned fish and from social information derived from local social interactions among group members.

Similar collective escape responses initiated by rapid changes in direction and speed have been documented in fish schools [6, 15], marine insects [38], and starling flocks [4, 23] during predator attacks. In these systems, only a small subset of individuals initially detects the threat, yet their behavioural response rapidly spreads through the group. For example, when sulphur mollies are attacked by birds, only the fish closest to the predator first perceive the threat and rapidly dive below the water surface [6]. Nearby individuals subsequently perform the same manoeuvre, generating propagating surface waves. As sulphur mollies rely on surface respiration, diving carries substantial physiological costs. It is therefore unlikely to occur in the absence of danger and consequently provides a highly reliable indicator of danger. Our results suggest that rapid increases in swimming speed play a similar signalling function in schooling fish, serving as socially transmitted indicators of risk.

The temporal structure of the threat may be crucial in shaping how private information spreads across groups. In systems such as starling flocks attacked by falcons [4], fish schools pursued by piscivorous predators [15], pigeon flocks escaping a robotic falcon [16], or African ungulate herds chased by lions [39], the threat persists throughout the escape, allowing multiple individuals to continuously acquire direct information about the predator while simultaneously responding to the movements of their neighbours. By contrast, our experiment involves a pulse-like perturbation: the green light constitutes an aversive stimulus only for the conditioned fish and only for a brief period. The collective response is then primarily maintained by internal social interactions after the initial perturbation has occurred. Similar pulse-like dynamics may occur during ambush-like or brief attacks, for example, a localised bird strike triggers a wave of diving fish [6]. Under such transient threats, the central challenge is to study the transmission of information about a short-lived threat available only to one or a few individuals. Our experiments show that despite the short-lived nature of threat information, the escape response is mediated entirely by internally propagated social information [40]. Apart from contexts in which predators continuously chase prey, our study also contrasts with migration or resource-tracking contexts, where informed individuals often possess relatively persistent directional information that can guide group movement over longer timescales [11, 12, 31, 13, 14, 21]. Thus, our study extends the framework for how private information transfers into social information from contexts of persistently available information to scenarios where information is local and short-lived.

By analysing temporal correlations in swimming speed, we identified evidence of information propagation from the conditioned fish to the rest of the group. Our results reveal the emergence of a hierarchical leadership structure aligned with the crossing order, such that the individual with crossing rank *i* most often follows the individual with crossing rank *i* − 1. This pattern is consistent with previous studies showing that faster-moving individuals tend to emerge as leaders in fish schools [41]. This sequential propagation of leadership subsequently extended to the remaining leader–follower pairs within the group. We further found that the maximum speed reached by the fish during hurdle crossing increased with crossing rank. As maintaining group cohesion is crucial for preserving the anti-predatory benefits of group living [42, 40], individuals remaining in the original compartment likely accelerate more strongly as an increasing number of group members have already moved to the opposite side of the tank. This progressive increase in speed reflects an attempt to maintain cohesion with the escaping subgroup. Thus, the group response emerges from the conditioned fish initiating escape based on private information, followed by a cohesion-maintaining response of the remaining naive fish mediated by speed matching. However, our conclusions are based on experiments conducted within the constrained box; whether the same mechanisms operate in open arenas or during natural predator attack remains to be tested.

Nevertheless, in about 30% of the trials, not all naive fish cross the hurdle. This observation suggests that social conformity effects [28] may sometimes inhibit the propagation of threat information. In particular, when the majority of group members remain inactive, the conditioned fish itself may be less likely to initiate escape, thereby preventing the spread of the response through the group. This reveals a fundamental trade-off between private information and social influence. An informed fish must respond strongly enough to initiate escape, while remaining partially socially coupled for its movement to recruit followers. While too much conformity may suppress escape initiation, complete social decoupling may reduce its ability to influence the rest of the group. This trade-off between private and social information may represent a rather general principle governing collective decision-making in animal groups.

Despite the sequential increase in speeds, we found no evidence of amplification of information during propagation. The latency between the onset of the stimulus and hurdle crossing increased linearly with the crossing rank, while the time interval between successive crossings remained nearly constant throughout the escape sequence. Similar sequential transmission without amplification has been reported during spontaneous collective U-turns in fish schools [28]. Thus, although fish progressively accelerated to restore cohesion with the escaping subgroup, each transmission step preserved approximately the same temporal characteristics. The propagation therefore resembles a relay mechanism rather than a positive-feedback cascade in which successive individuals would respond increasingly rapidly. The absence of information amplification is also consistent with the idea that each fish interacts with only a limited subset of its neighbours, rather than integrating information from the entire group [43, 26, 44]. By restricting social attention to a few influential neighbours, individuals effectively filter incoming information. This allows salient behavioural perturbations to propagate while preventing the amplification of behavioural fluctuations (including noise) and limiting the spread of false signals. Similar selective information processing has been documented in several collective systems, including fish schools, bird flocks, and insect societies, where individuals dynamically weight only the most relevant social cues available in their local neighbourhood [26, 41].

Finally, using a computational model, we demonstrated that local social interaction rules are sufficient to reproduce the main collective properties observed experimentally. In particular, the model successfully captured the emergence of hierarchical leadership dynamics based solely on local velocity-matching and distance-regulation interactions, along with a temporary reduction in the strength of social interaction of the conditioned fish following threat perception. Importantly, the model revealed that conditioned individuals cannot fully decouple socially from their neighbours during escape. Residual social coupling proved necessary to reproduce the experimentally observed propagation dynamics, suggesting that informed individuals remain partially embedded within collective decision-making even as they respond to threats. Our findings also emphasise the importance of incorporating variable-speed dynamics together with context-dependent interaction rules into models of collective escape [37, 22, 45, 41, 46]. Consequently, the constant-speed assumption commonly adopted in classical schooling and flocking models may overlook key mechanisms that underlie the propagation of escape responses in animal groups.

In summary, our study demonstrates that aversive conditioning provides a powerful experimental framework for precisely controlling the source of threat information within an animal group and for studying the mechanisms by which an initially private response leads to a collective escape. These results suggest that collective escape emerges when local responses by informed individuals are transmitted through relatively simple local interactions among limited numbers of neighbours. Future work will be required to reconstruct the detailed interaction rules governing escape behaviour at the individual level [47]. Furthermore, our framework of aversive conditioning could be used to design experiments that control not only informed individuals, but also the number of informed individuals, how long information persists, and whether different individuals receive conflicting cues. Such experiments are essential for understanding collective responses to coordinated predator attacks, conflicting information, and complex collective decision-making scenarios [15, 48].

## Ethics statement

All experimental procedures were approved by the Institutional Animal Ethics Committee at the Indian Institute of Science, Bengaluru, India (CAF/Ethics/851/2021). Permits were also obtained from the National Biodiversity Authority of the Government of India (INBA1202203442). All procedures were designed to minimise stress and handling. No animals were sacrificed during this study.

## Acknowledgements

VJ, CA, GT and VG acknowledge support from the Indo-French Center for the Promotion of Advanced Research (Project No. 64T4-B). CA was also supported by an IoE postdoctoral fellowship from the Indian Institute of Science, and VJ from the Prime Minister’s Research Fellowship program (Ministry of Education, Government of India). DT was funded by the European Union’s Horizon 2020 research and innovation program under the Marie Skłodowska-Curie grant agreement No. 101154645. GT also gratefully acknowledges the Indian Institute of Science for support via the Infosys Visiting Chair Professor at the Centre for Ecological Sciences, IISc, Bengaluru. We thank Maud Combe for developing the custom programs used for data analysis and Anshi Pillai for helping with tracking. We thank Wenying Shou and Alex E. Yuan for providing insights into the permutation method. We thank Hitesh C K for comments on the code.

