## Supplementary material for "Tracing information propagation in fish schools with a conditioned escape response": SM_bioRxiv.pdf

August 4, 2026

**This PDF file includes:**

- **Supplementary Text**
- **Figures S1-S11**
- **Table S1-S2**

### Supplementary Text

#### S1 Experimental setup and procedure

##### S1.1 Study animals

We conducted an avoidance conditioning protocol on the tiger barb, *Puntigrus tetrazona*, a tropical freshwater schooling species. Seventy males ($3.73 \pm 0.18$ cm) were obtained from the Fisheries Research & Information Centre (FRIC) in Hesaraghatta, Bengaluru, India. Upon arrival, the fish were transferred to a 160 L holding tank ($89\times45\times40\,\mathrm{cm}^3$) filled with aged water consisting of 70% reverse-osmosis (RO) and 30% tap water. The water was continuously aerated, maintained at $27 \pm 0.5$°C, and partially renewed (20%) once a week. The fish were fed *ad libitum* with commercial food pellets (Optimum$^{\mathrm{TM}}$). Ambient lighting was automatically controlled to maintain a 12 h:12 h light–dark cycle. Fish were acclimated to laboratory conditions for 15 days before the start of the experiments.

##### S1.2 Conditioning procedure

The aversive conditioning protocol was adapted from Lecheval and Theraulaz [1]. One week after their arrival at the facility, 14 fish were randomly selected from the holding tanks and tagged with Visible Implant Elastomer at the base of the dorsal fin (Northwest Marine Technology, NMT Inc, Shaw Island, WA) following the manufacturer's instructions and subsequently housed individually. These fourteen fish were subjected to conditioning (hereafter referred to as conditioned individuals).

Conditioning was conducted in a shuttle box ($50\times20\times20\,\mathrm{cm}^3$) consisting of two compartments ($20\times20\times20\,\mathrm{cm}^3$) separated by a 7.5 cm-high and 10 cm-wide trapezoidal hurdle positioned in the centre. Each compartment was equipped with 84 green light-emitting diodes (LEDs), arranged in four rows of seven on three sides of the tank, which could be controlled independently (figure 1a,b). Two stainless steel mesh electrodes in each compartment, connected to a stimulator, delivered mild, controlled electric shocks. The current was set to the minimum intensity to elicit a visible behavioural response from the fish (7 V, 2.7 mA measured at the electrodes). Both visual and electrical stimuli were delivered manually. The box was filled with freshwater of controlled composition (70% RO and 30% tap) to a depth of 10 cm (2.5 cm above the hurdle). The experimental setup was homogeneously illuminated and surrounded by an opaque white curtain to minimise external visual disturbances.

Conditioning sessions were conducted over a period of 40 days (22 sessions per fish). Prior to each session, conditioned fish were placed in the shuttle box and allowed to acclimate for 10 min. Each session consisted of 10 trials separated by a 2 min inter-trial interval. The water was renewed after each session. Training sessions were recorded using a digital camera (Canon EOS-600D) positioned 1 m above the box, with a spatial resolution of 1920×1080 pixels and a frame rate of 25 frames per second.

During each trial, the conditioned stimulus (CS; green light) was presented for 3 sec in the compartment occupied by the focal fish. If the fish failed to cross the hurdle within this period, the unconditioned stimulus (US; electric shock) was delivered. A trial was considered successful if the fish crossed the hurdle before the onset of the US (i.e., within 3 sec of the onset of the CS; figure 1c). The escape performance was quantified as the proportion of successful trials out of 10. Fish underwent repeated conditioning until reaching a performance criterion of 80% (i.e., at least 8 successful trials out of 10 without receiving the US). Only individuals meeting this criterion were used in subsequent collective escape experiments (7 out of 14 fish). After the conditioning session, fish were returned to the holding tank.

##### S1.3 Experimental protocol

To investigate the collective escape response, groups composed of one conditioned and four naive fish were assembled 24 h prior to testing. Individuals were transferred from the holding tanks to an intermediate tank (26×29×59 $cm^3$) containing water with the same properties as the holding conditions. This intermediate step was used to facilitate acclimation and to minimise handling stress on the day of the experiment. The shuttle box was filled with freshwater (70% RO and 30% tap) at 27°C to a depth of 10 cm. Fish were transferred from the intermediate tank to the shuttle box and allowed to acclimate for 10 min before the test began. Each trial consisted of 3 second presentation of the conditioned stimulus (green light), during which collective behaviour was recorded. Trials were separated by a 2 min inter-trial interval (figure 1d). Trials were recorded at 100 frames per second (Sony PXW-FS5). The order in which the conditioned fish were tested with the naive fish was randomised. After each session, conditioned fish were returned to their individual holding tanks, whereas naive fish were transferred to a separate post-experimental tank. The water in the shuttle box was replaced between sessions.

As a control experiment, 6 groups of 5 naive fish were tested in the shuttle box in the absence of electric shocks to quantify the baseline probability of escape responses induced by the green light alone.

##### S1.4 Tracking and processing

Fish trajectories were extracted using a custom tracking software specifically developed at CRCA. Individual fish were tracked using a graphical user interface that enabled manual validation and correction of trajectories.

##### S1.5 Quantification of collective escape response

We denote the raw time in seconds by $\tilde{t}$, and define $\tilde{t}_0$ as the onset of the green light and $\tilde{t}_{\max}$ as the time at which the last fish completely crosses the hurdle. Then, we introduce a normalised time $t$ so that the escape phase spans $[0,1]$ in all trials:

$$t = \frac{\tilde{t} - \tilde{t}_0}{\tilde{t}_{\max} - \tilde{t}_0}. \quad (1)$$

Accordingly, $t < 0$ corresponds to the initial phase, whereas $t > 1$ corresponds to the relaxation phase. This normalisation enables direct comparison of escape dynamics across trials.

The position and velocity vectors of fish $i$ are denoted by $\vec{r}_i\,(x_i, y_i)$, $\vec{v}_i\,\left(v_x^i, v_y^i\right)$. The position of the barycentre $B$ is given by $\vec{r}_B = \frac{1}{N}\sum_{i=1}^{N} \vec{r}_i$, where $N$ is the total number of fish.

The collective behaviour is quantified using two standard metrics:

- *Group dispersion* quantifies the spatial cohesion of the group and is defined as the mean distance of individuals from the group barycentre $B$:

$$D(t) = \frac{1}{N}\sum_{i=1}^{N} d_{Bi}(t), \quad D_x(t) = \frac{1}{N}\sum_{i=1}^{N} d_{Bi}^x(t), \quad D_y(t) = \frac{1}{N}\sum_{i=1}^{N} d_{Bi}^y(t), \quad (2)$$

  where $d_{Bi} = \|\vec{r}_i - \vec{r}_B\|$ is the distance between $i$ and $B$, with $d_{Bi}^x = \|x_i - x_B\|$ along $x$-axis, and similar expression in the $y$-coordinate. We also define the dispersion $D_{\rm wc}$, computed *without the conditioned* fish, after excluding $i = 1$ from the sums in (2).

- *Group polarisation* quantifies the degree of alignment among fish:

$$P(t) = \frac{1}{N}\left\|\sum_{i=1}^{N} \frac{\vec{v}_i(t)}{\mathrm{v}_i(t)}\right\|, \quad P_x(t) = \frac{1}{N}\left|\sum_{i=1}^{N} \frac{v_i^x(t)}{\mathrm{v}_i(t)}\right|, \quad P_y(t) = \frac{1}{N}\left|\sum_{i=1}^{N} \frac{v_i^y(t)}{\mathrm{v}_i(t)}\right|, \quad (3)$$

  where $\mathrm{v}_i = \|\vec{v}_i\|$ is the speed of individual $i$.

#### S2 Modelling collective escape dynamics

**Model rules:** The motion of fish $i$, conditioned or naive, is governed by

$$\frac{d\mathrm{v}_i}{dt}(t) = \vec{F}_i(t) \cdot \vec{e}^{\,i}_{\parallel}(t), \qquad \frac{d\phi_i}{dt}(t) = \frac{1}{\mathrm{v}_i(t) + \alpha}\, \vec{F}_i(t) \cdot \vec{e}^{\,i}_{\perp}(t), \tag{4}$$

where $\mathrm{v}_i$ and $\phi_i$ denote the speed and the heading angle of the fish, $\vec{e}^{\,i}_{\parallel}$ and $\vec{e}^{\,i}_{\perp}$ are unit vectors respectively parallel and perpendicular to the direction of motion of the fish, $\alpha$ is a rotational friction coefficient, and $\vec{F}_i$ is the total force acting on the fish, which depends on whether it is the conditioned one or not:

$$\vec{F}_i(t) = \begin{cases} \vec{F}^i_{\mathrm{sp}}(t) + \vec{F}^i_{\mathrm{Ali}}(t) + \vec{F}^i_{\mathrm{Att}}(t) + \vec{F}^i_{\mathrm{w}}(t) & \text{for naive fish,} \\ \vec{F}^c_{\mathrm{sp}}(t) + (1-\omega(t)) \left[\vec{F}^c_{\mathrm{Ali}}(t) + \vec{F}^c_{\mathrm{Att}}(t) + \vec{F}^c_{\mathrm{w}}(t)\right] + \omega(t)\vec{F}_{\mathrm{esc}}(t) & \text{for the conditioned fish.} \end{cases}$$

where we will define each of the force terms below.

The self-propulsion $\vec{F}^i_{\mathrm{sp}}(t)$ makes fish adopt a preferred speed $\mathrm{v}_0$ with relaxation rate $\beta$, and includes independent Gaussian noises accounting for fish swimming fluctuations,

$$\vec{F}^i_{\mathrm{sp}}(t) = \beta\big(\mathrm{v}_0 - \mathrm{v}_i(t)\big)\, \vec{e}^{\,i}_{\parallel}(t) + \sqrt{2D_{\mathrm{v}}}\, \eta_{\mathrm{v}}(t)\, \vec{e}^{\,i}_{\parallel}(t) + \sqrt{2D_{\phi}}\, \eta_{\phi}(t)\, \vec{e}^{\,i}_{\perp}(t), \tag{5}$$

where $D_{\mathrm{v}}$ and $D_{\phi}$ are the intensity of speed and heading fluctuations, respectively.

Social interactions are described by a force of alignment $\vec{F}^i_{\mathrm{Ali}}(t)$ and a distance-regulation force $\vec{F}^i_{\mathrm{Att}}(t)$ with neighbours. We consider that fish pay attention to two neighbours $j_1$, $j_2$ chosen randomly in the visual field of fish $i$ [2–4]. For small groups ($N = 5$), this neighbourhood rule is consistent with empirical observations [5, 6], although the model's qualitative behaviour remains robust under alternative neighbourhood definitions, provided the interaction strengths are appropriately recalibrated. Thus,

$$\vec{F}^i_{\mathrm{Ali}}(t) = \frac{\mu_{\mathrm{Ali}}}{2}\left[\vec{v}_{j_1}(t) + \vec{v}_{j_2}(t) - 2\vec{v}_i(t)\right], \tag{6}$$

$$\vec{F}^i_{\mathrm{Att}}(t) = \frac{\mu_{\mathrm{Att}}}{2} \sum_{j_1, j_2} \tanh\left[2.5(r_{ji}(t) - r_D)\right] \vec{e}_{ji}(t), \tag{7}$$

where $r_{ji} = \|\vec{r}_j - \vec{r}_i\|$, $\vec{e}_{ji}$ is the unit vector pointing from $i$ to $j$, and $r_D$ is a fixed preferred inter-individual distance. The sign of $r_{ji} - r_D$ determines the attractive/repulsive character of $\vec{F}^i_{\mathrm{Att}}$.

The effect of the force $\vec{F}^i_{\mathrm{w}}(t)$ exerted by a tank wall is to align fish velocity with the wall, with an intensity that decays exponentially with the distance to it, simulating a soft rebound:

$$\vec{F}^i_{\mathrm{w}}(t) = \mu_{\mathrm{w}} \exp\left(-\frac{r^i_{\mathrm{w}}(t)}{l_{\mathrm{w}}}\right) \left(\vec{e}^{\,i}_{\mathrm{w}}(t) - \vec{e}^{\,i}_{\parallel}(t)\right), \tag{8}$$

where $r_{\mathrm{w}}^i(t)$ is the distance to the nearest wall, $l_{\mathrm{w}}$ is the spatial range of interaction, and $\vec{e}_{\mathrm{w}}^{\,i}(t)$ is the unit vector parallel to the nearest wall, oriented in the same half-plane as the velocity vector of fish $i$. Moreover, the force is applied only when the fish is moving towards the wall. When simulating the initial and relaxation phases, both the boundaries of the tank and the central hurdle are treated as walls and contribute to $\vec{F}_{\mathrm{w}}^{i}(t)$ through equation (8). When the green light is turned on, the hurdle no longer exists for the fish.

For the conditioned fish, the total force includes an additional term $\vec{F}_{\mathrm{esc}}(t)$ which is activated when the green light is turned on, via a modulation factor, $\omega(t) \in [0, 1]$; this force term captures the aversion of the conditioned fish to the green light, modelled as attraction towards the centre of the adjacent compartment $\vec{R}$ and is given by

$$\vec{F}_{\mathrm{esc}}(t) = \mu_{\mathrm{esc}}\, \vec{e}_R(t), \tag{9}$$

where $\vec{e}_R(t) = (\vec{R} - \vec{r}_c(t))/\|\vec{R} - \vec{r}_c(t)\|$ is the unit vector pointing from the conditioned fish to $\vec{R}$, and $\mu_{\mathrm{esc}}$ is the maximum intensity of the escape force. Note that $\omega(t) = 0$ for $t < 0$, and is set to a value of $\omega_0$ from the time the light is turned on to the time the conditioned fish crosses the hurdle, after which it decays following $\omega(t) = \omega_0 e^{-\gamma t}$.

Simulations were run with boundary conditions that mimic the experimental geometry: a rectangular domain of $50{\times}20\,\mathrm{cm}^2$, divided into two $20{\times}20\,\mathrm{cm}^2$ compartments separated by a central, 10 cm-wide rectangle representing the hurdle. We set the preferred speed and preferred inter-individual distance to values comparable to empirical measurements. Social interaction strengths were selected such that simulated individuals exhibited swimming speeds comparable to those observed experimentally during the initial and relaxation phases. We varied the modulation factor of the escape force $\omega(0)$ to investigate its influence on the collective properties observed during escape events. Detailed parameter values are given in the table S2.

To understand the implications of social interactions and the influence of geometric constraints, we consider a null model by setting $\mu_{\mathrm{Ali}} = 0$ and $\mu_{\mathrm{Att}} = 0$. In this null model, when the light is turned on, all fish behave like conditioned fish (section S3).

#### S3 Null model

We developed a null model in which individuals do not interact socially with neighbours but instead respond independently to the external stimulus. As in the model described in Section 3c, all fish align with the tank walls. At the onset of the green light stimulus ($t = 0$), each fish moves toward the centre of the adjacent compartment according to the escape dynamics described in Section 3c. Because all individuals respond directly to the stimulus in this null model, there is no predefined conditioned fish; instead, the first fish to cross the hurdle is retrospectively designated as the conditioned fish. Simulations were performed in a tank matching the dimensions of the experimental setup. The null model therefore provides a baseline for distinguishing collective responses emerging from social interactions from those resulting solely from independent responses to the external stimulus.

We found that group polarisation increased in a manner qualitatively similar to the experimental observations. This indicates that this pattern largely arises from the geometric constraints imposed by the experimental tank, in which fish movement is effectively restricted to a single spatial axis during escape. However, unlike the empirical observations, the null model predicted that the fish closest to the conditioned fish crossed the hurdle immediately afterward. More generally, fish cross the hurdle primarily according to their initial proximity to the hurdle. Finally, in the null model, the time interval between successive crossings increased with crossing rank, whereas in the empirical data, this interval either decreased or remained constant. These differences indicate that the experimentally observed escape dynamics cannot be explained solely by independent responses to the stimulus and instead require socially mediated interactions between individuals.

#### S4 Crossing rank and initial spatial configuration

To investigate whether crossing order was related to the initial spatial configuration of the group, we quantified the distance between each naive fish and the conditioned individual at $t = 0$. Let $\vec{r}_C(0)$ and $\vec{r}_i(0)$ denote position vectors of the conditioned fish and the naive fish with crossing rank $i$, respectively. The Euclidean distance and longitudinal distance between the conditioned fish and naive fish $i$ were defined as $r^i_C = \|\vec{r}_C(0) - \vec{r}_i(0)\|$ and $x^i_C = |x_C(0) - x_i(0)|$, respectively. In addition, at $t = 0$, we measured the viewing angle $\psi_{iC}$, defined as the angular deviation required for naive fish $i$ to orient toward the conditioned fish, and the relative orientation, $\phi_{iC}$, defined as the difference in heading angle between naive fish $i$ and conditioned fish. By convention, angles are positive for counterclockwise rotations and negative for clockwise rotations. These measurements were used to test whether crossing rank was associated with the initial spatial arrangement and orientation of individuals relative to the conditioned fish.

We found no significant relationship between crossing rank and the initial distance to the conditioned fish (Euclidean distance, LME: $\chi^2(3) = 1.99$, $p = 0.57$, figure S4a). Likewise, neither the viewing angle ($\psi_{iC}$) nor the relative orientation ($\phi_{iC}$) had a significant effect on the crossing rank (LME: $\chi^2(3) = 0.42$, $p = 0.93$, figure S4b; LME: $\chi^2(3) = 1.33$, $p = 0.72$, figure S4c). These results indicate that the initial spatial position and orientation of naive individuals relative to the conditioned fish do not predict the order in which they cross the hurdle during the collective escape response.

#### S5 Leader-follower relationships and information propagation

To quantify leader–follower relationships during escape propagation, we compute the cross-correlation of individual swimming speeds, which measures the degree to which speed fluctuations are copied after a time delay $\tau$. Given the speed time series $\mathrm{v}_i(t)$ and $\mathrm{v}_j(t)$ of fish $i$ and $j$, respectively, the normalised cross-correlation at lag $\tau$ is defined as:

$$C(\mathrm{v}_i, \mathrm{v}_j; \tau) = \frac{1}{T\sigma_i\,\sigma_j} \sum_{t=1}^{T-\tau} \big(\mathrm{v}_i(t+\tau) - \bar{\mathrm{v}}_i\big)\big(\mathrm{v}_j(t) - \bar{\mathrm{v}}_j\big), \qquad \text{(S1)}$$

where $\bar{\mathrm{v}}_i$, $\bar{\mathrm{v}}_j$ and $\sigma_i$, $\sigma_j$ are the mean and standard deviation of each speed time series computed over the time window ($t = 1, \ldots, T$) of an escape phase.

We then define the peak lag $\tau_{ij}$ as the value of $\tau$ at which $C(\mathrm{v}_i, \mathrm{v}_j; \tau)$ reaches its maximum. Individual $i$ is considered to lead $j$ when $\tau_{ij} < 0$, whereas $j$ is considered to lead $i$ when $\tau_{ij} > 0$. The values $\tau_{ij}$ for all pairs $(i, j)$ allow us to construct a leader-follower network for each trial. This method, particularly when applied to directional correlations, has been widely used to investigate leader-follower dynamics in collectively moving groups under both perturbed [7] and unperturbed conditions [8–10].

However, the temporal correlation function defined in equation (S1) is strictly applicable only to stationary time series (figure S7). While the assumption of stationarity may hold during the initial and relaxation phases of our experiments, the speed time series recorded during the escape phase are clearly non-stationary, being characterised by rapid accelerations followed by decelerations. Although several approaches have been proposed to infer correlations from non-stationary signals – such as detrending the data over selected temporal windows to recover approximate stationarity – these methods remain highly sensitive to methodological assumptions, including the choice of window size [11]. This sensitivity substantially limits the robustness and interpretability of the inferred correlations.

Alternatively, nonparametric approaches, such as permutation tests, can be used to reliably assess the significance of correlations in non-stationary time series [12, 13]. The principle underlying these methods is that, if two variables are genuinely correlated, correlations computed within the same trial should be significantly stronger than correlations computed between independent trials. By analysing multiple independent and identically distributed trials, the significance of within-trial correlations can therefore be evaluated relative to a null distribution constructed from between-trial correlations [14, 15].

For example, let $\mathrm{v}_1^k$ and $\mathrm{v}_2^k$ denote the speed time series of the fish with crossing rank 1 (the conditioned fish) and crossing rank 2 (the first fish to cross the hurdle after the conditioned fish) during trial $k$. The within-trial correlation $C(\mathrm{v}_i^k, \mathrm{v}_j^k; \tau_{i^k j^k})$ can then be compared with surrogate correlations of the form $C(\mathrm{v}_i^k, \mathrm{v}_j^l; \tau_{i^k j^l})$ where $l \neq k$. This procedure, commonly referred to as inter-subject surrogates method, provides a nonparametric framework for evaluating the significance of correlations between two fish within a trial. In practice, surrogate correlations are computed for all $C(\mathrm{v}_i^k, \mathrm{v}_j^l; \tau_{i^k j^l})$ for $l = 1, 2, 3, \ldots, O$, where $O$ denotes the total number of trials. The associated p-value was defined as the fraction of surrogate correlations exceeding the original within-trial correlation, $C(\mathrm{v}_i^k, \mathrm{v}_j^k; \tau_{i^k j^k})$. Because the total number of trials was relatively limited in our dataset ($O = 53$), this approach imposes a lower bound on the attainable significance level ($1/O = 0.0189$). However, since our primary objective was to characterise the overall propagation of information from conditioned to naive fish, we additionally performed a global permutation test pooling data across all trials [16–18].

To illustrate this procedure using three trials as an example, we first compute the mean within-trial correlation between $\mathrm{v}_1$ and $\mathrm{v}_2$ across trials $k = 1, 2, 3$:

$$C(\mathrm{v}_1, \mathrm{v}_2) = \frac{1}{3}\left[C(\mathrm{v}_1^1, \mathrm{v}_2^1) + C(\mathrm{v}_1^2, \mathrm{v}_2^2) + C(\mathrm{v}_1^3, \mathrm{v}_2^3)\right]. \tag{S2}$$

Next, surrogate datasets were generated by randomly shuffling the $\mathrm{v}_2$ time series across trials while preserving the original ordering of $\mathrm{v}_1$. One example of such a permutation is $k = 2, 3, 1$:

$$C_{\mathrm{shfl}}(\mathrm{v}_1, \mathrm{v}_2) = \frac{1}{3}\left[C(\mathrm{v}_1^1, \mathrm{v}_2^2) + C(\mathrm{v}_1^2, \mathrm{v}_2^3) + C(\mathrm{v}_1^3, \mathrm{v}_2^1)\right]. \tag{S3}$$

Correlations were then computed for all shuffled realisations. The significance of the original within-trial correlation was assessed as the proportion of shuffled correlations exceeding the original mean correlation ($C(\mathrm{v}_1, \mathrm{v}_2)$). Thus, the correlation between the speeds of fish with crossing ranks 1 and 2, given by equation (S2), was considered significant when fewer than 5% of shuffled realisations produced correlations larger than the observed original value ($p < 0.05$). Because the total number of trials was 53, exhaustive enumeration of all possible permutations (53!) was computationally intractable. We generated 10 000 random permutations to estimate the null distribution.

In addition to the strength of the correlation, we are interested in the leader-follower relationship determined via temporal lag. We note that bidirectional links between two individuals may occur; i.e., fish with crossing rank $i$ may significantly influence fish with crossing rank $j$ in some trials, whereas in others the influence may be reversed, such that both directional correlations are statistically significant. After identifying all significant leader-follower pairs, we constructed a leadership network in which each node represents a fish characterised by its crossing rank, and directed edges are drawn from followers to leaders.

##### A note on constructing the leader-follower network in the collective escape model

To construct the leadership network in the model, we followed the same method proposed in section S5. Here, we calculated the significance of correlations using a permutation test based on a sample size equivalent to our experimental trials ($N = 53$). To verify consistency, we then performed 100 independent sets of such simulations, each consisting of 53 trials.

#### Supplementary Figures

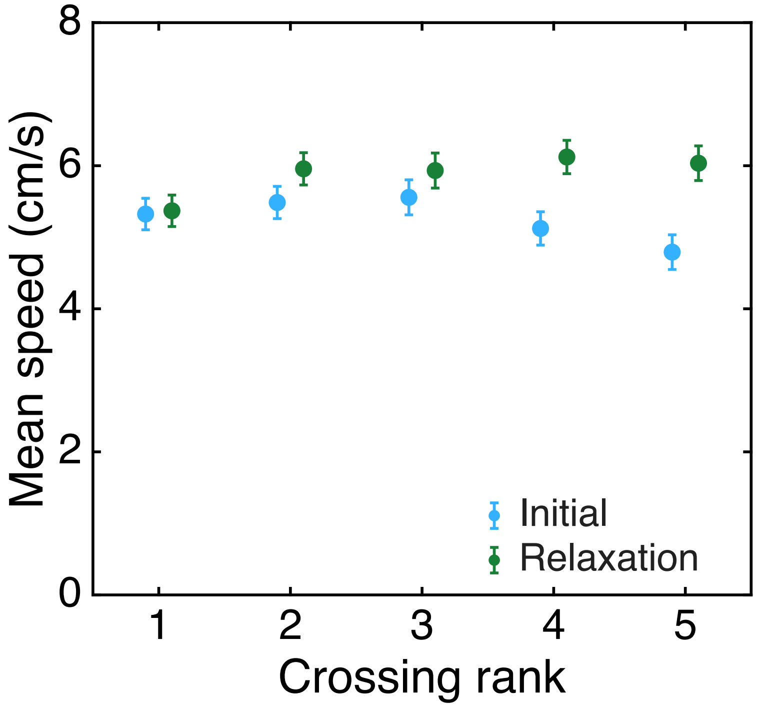


**Figure S1. Baseline locomotor behaviour of conditioned and naive fish.** Comparison of the mean swimming speed of the five fish (conditioned + naive fish) during the initial (blue) and relaxation (green) phases. These analyses were performed to assess whether aversive conditioning altered baseline locomotor behaviour. Conditioned and naive fish exhibited similar swimming speeds during both phases. We found no significant relationship between swimming speed and crossing rank during the initial phase (LME: $\chi^2(4) = 5.67$, $p = 0.23$). However, we found an overall significant relationship between swimming speed and crossing rank in the relaxation phase (LME: $\chi^2(4) = 54.09$, $p < 0.001$). Pairwise comparisons show that while CR-1 is slower ($5.4 \pm 0.24$ cm/s) than all subsequent fish, no significant differences in swimming speed were observed among fish with crossing rank 2–5 (with the maximum speed observed for CR-4, $6.12 \pm 0.24$ cm/s). Although we observe a statistical difference between conditioned and naive fish during the relaxation phase, the magnitude of this difference is very small ($\lesssim 0.2$ body length/s).

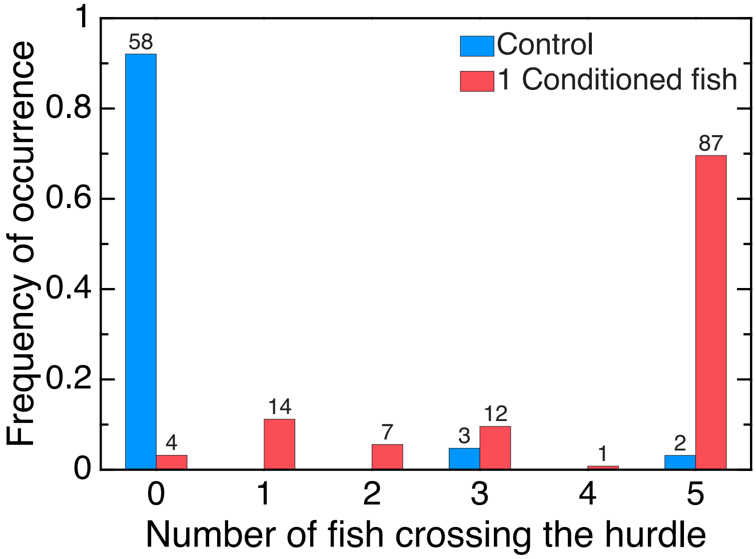


**Figure S2. Collective response to the conditioned stimulus in control and mixed groups ($N = 5$).** Response of control groups (blue) and groups containing one conditioned fish (red) to the green light stimulus. The numerical values at the top of each bar represent the number of trials with a given number of fish crossing the hurdle.

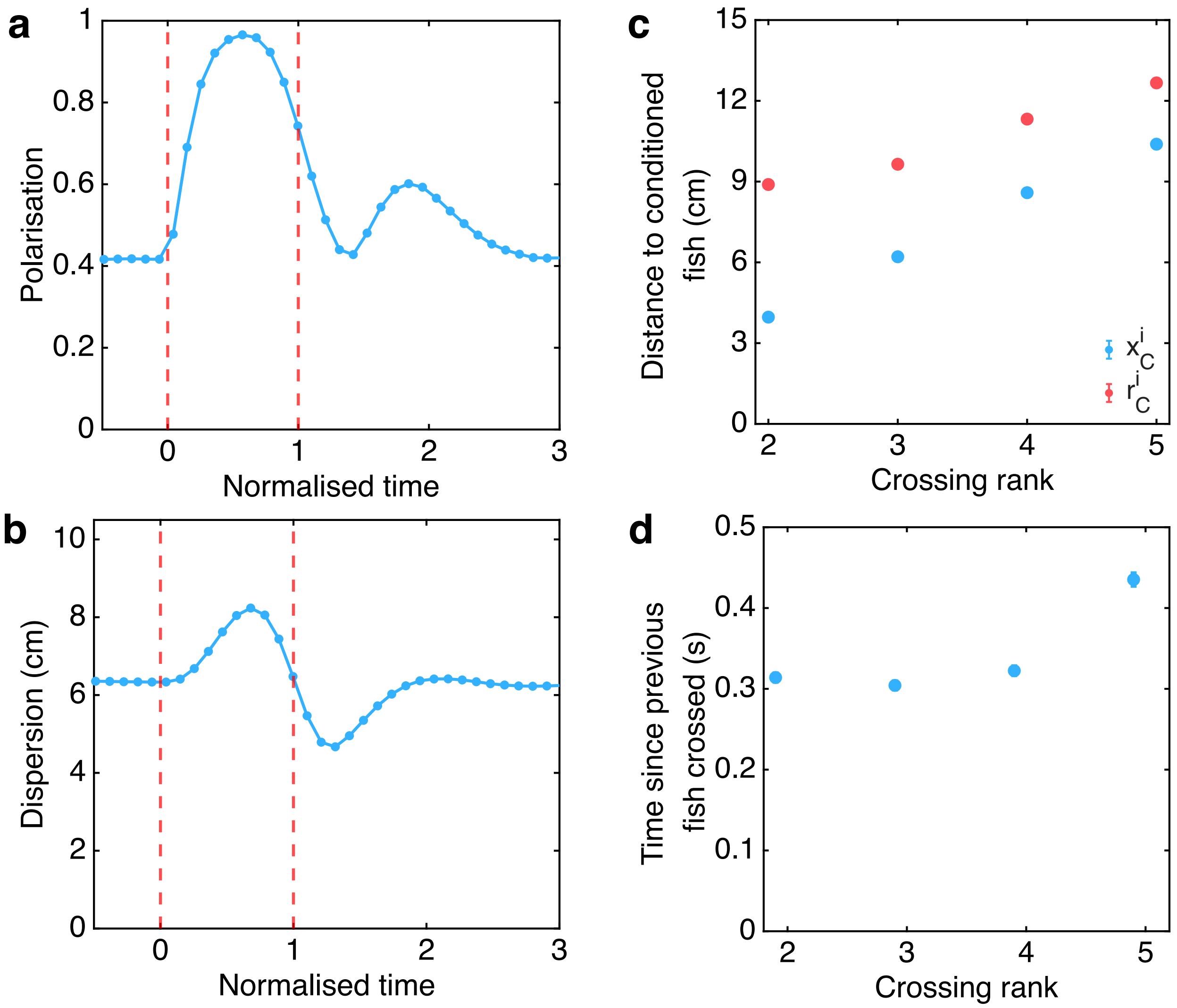


**Figure S3. Null-model dynamics of collective escape.** Average time series of (**a**) group polarisation and (**b**) group dispersion. The model reproduces the increase in polarisation during the escape phase arising from the geometric constraints of the experimental setup. (**c**) Relationship between crossing rank and initial distance to the conditioned fish measured using both longitudinal (blue) and Euclidean (red) distances. (**d**) Mean time interval between successive hurdle crossings as a function of crossing rank.

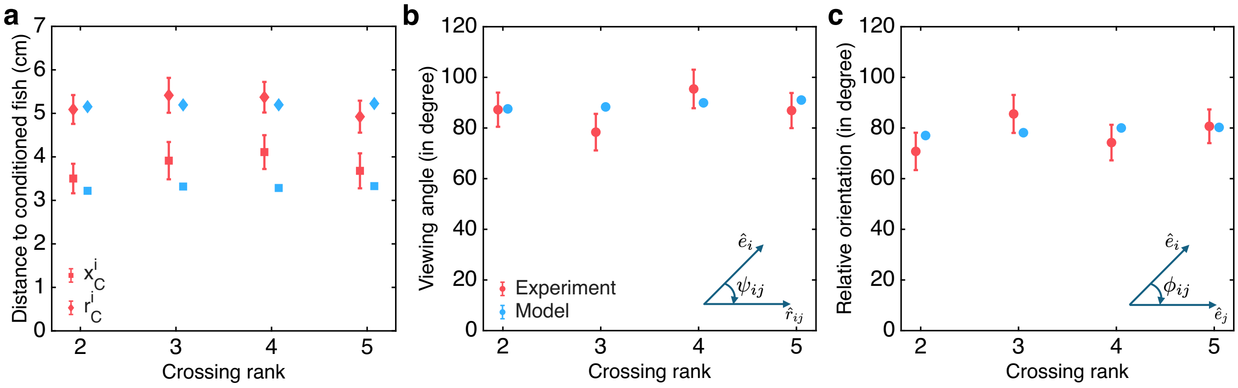


**Figure S4. Relationship between crossing rank and the initial spatial configuration relative to the conditioned fish at the onset of green light.** (**a**) Crossing rank is not associated with the initial distance between naive individuals and the conditioned fish, whether measured as Euclidean distance (diamond) and longitudinal distance along the x-axis (square). (**b**) No relationship is observed between crossing rank and the viewing angle of the naive fish relative to the conditioned fish. Inset: Schematic representation of the viewing angle ($\psi_{ij}$). (**c**) Crossing rank is also unrelated to the relative orientation between the naive and conditioned fish. Inset: Schematic representation of the relative orientation ($\phi_{ij}$). Blue symbols correspond to experimental data, whereas red symbols indicate results from the computational model. Error bars represent the standard error.

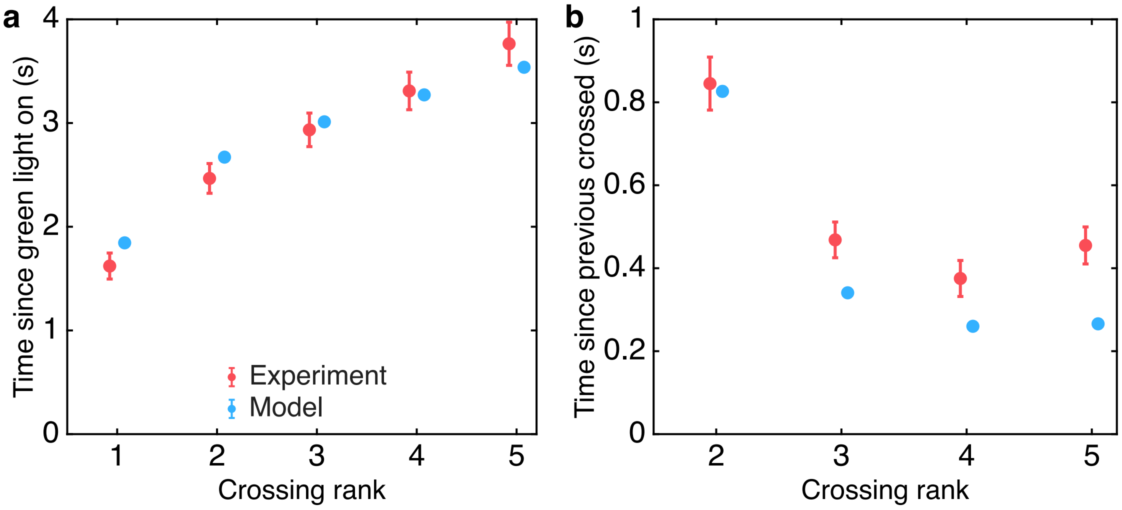


**Figure S5. Sequential propagation of collective escape response.** (a) Mean latency to cross the start of the hurdle as a function of crossing rank measured from the onset of the green light stimulus. Crossing latency increases approximately linearly with crossing rank. (b) Mean time interval between successive crossings. The first naive fish (crossing rank 2) crosses the start of the hurdle approximately 0.8 s after the conditioned fish. The inter-crossing interval decreases to $\approx 0.4$ s between crossing ranks 2 and 3 and remains relatively constant for subsequent crossings. Red symbols correspond to experimental data, and blue symbols represent results from the computational model. Error bars indicate the standard error.

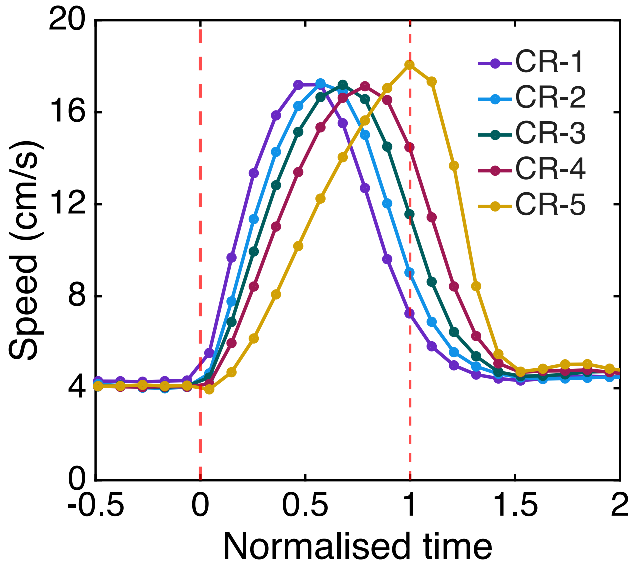


**Figure S6. Time series of individual speeds sorted by crossing rank for the null model.** As each individual responds independently to the perturbation, speed during the escape phase is clearly non-stationary, characterised by rapid accelerations followed by decelerations. Consequently, naively constructing a leader-follower network using the speed time series from the null model (equation (S1)) yields a spurious hierarchical network.

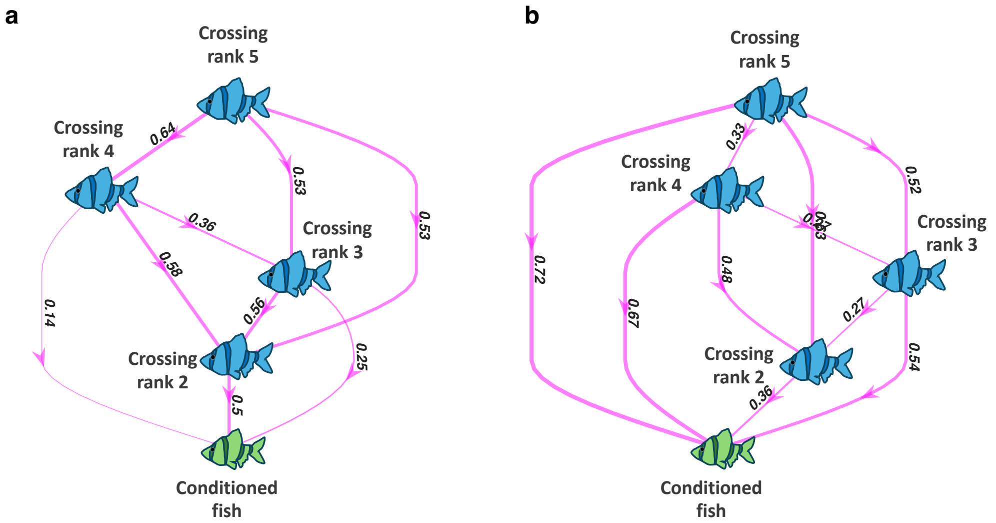


**Figure S7. Leadership network inferred from temporal speed correlations.** Leader–follower networks reconstructed using the correlation function defined in equation (S1) for (**a**) the experimental data and (**b**) the null model. Fish are labelled according to their crossing ranks, and directed edges are drawn from followers to leaders. Notably, the null model, despite lacking social interactions, also produces edges oriented from lower to higher crossing ranks, similarly to the experimental data. This apparent structure arises because the speed time series are intrinsically non-stationary, whereas the correlation function defined in equation (S1) assumes stationarity.

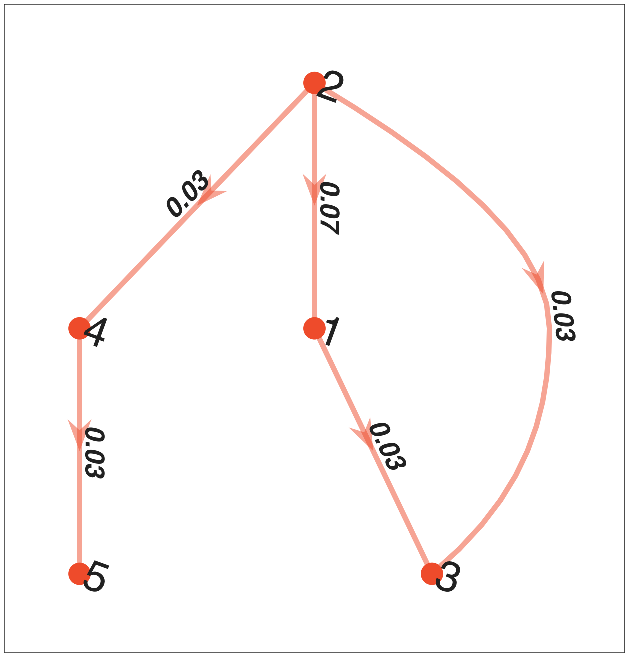


**Figure S8. Leadership network inferred from temporal speed correlations using the permutation method for the null model.** Fish are labelled according to their crossing ranks, and directed edges point from followers to leaders. Edge labels indicate the fraction of simulations in which the pairwise correlation between the two nodes was statistically significant. As expected, this method correctly infers no interaction between individuals in the null model.

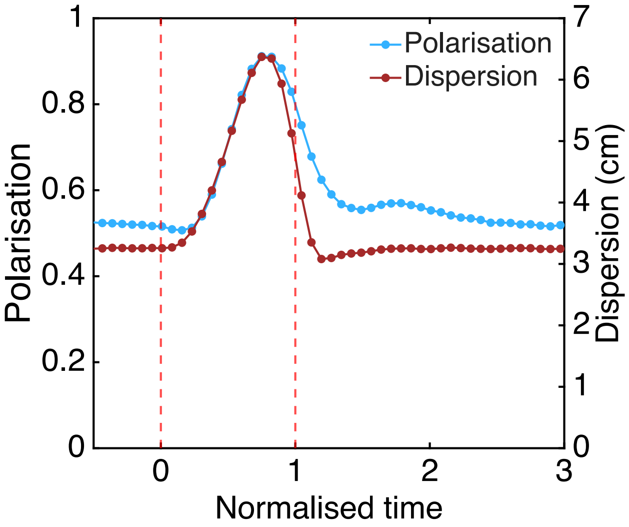


**Figure S9. Simulation results from the computational collective escape model.** Average time series of polarisation (blue) and dispersion (brown).

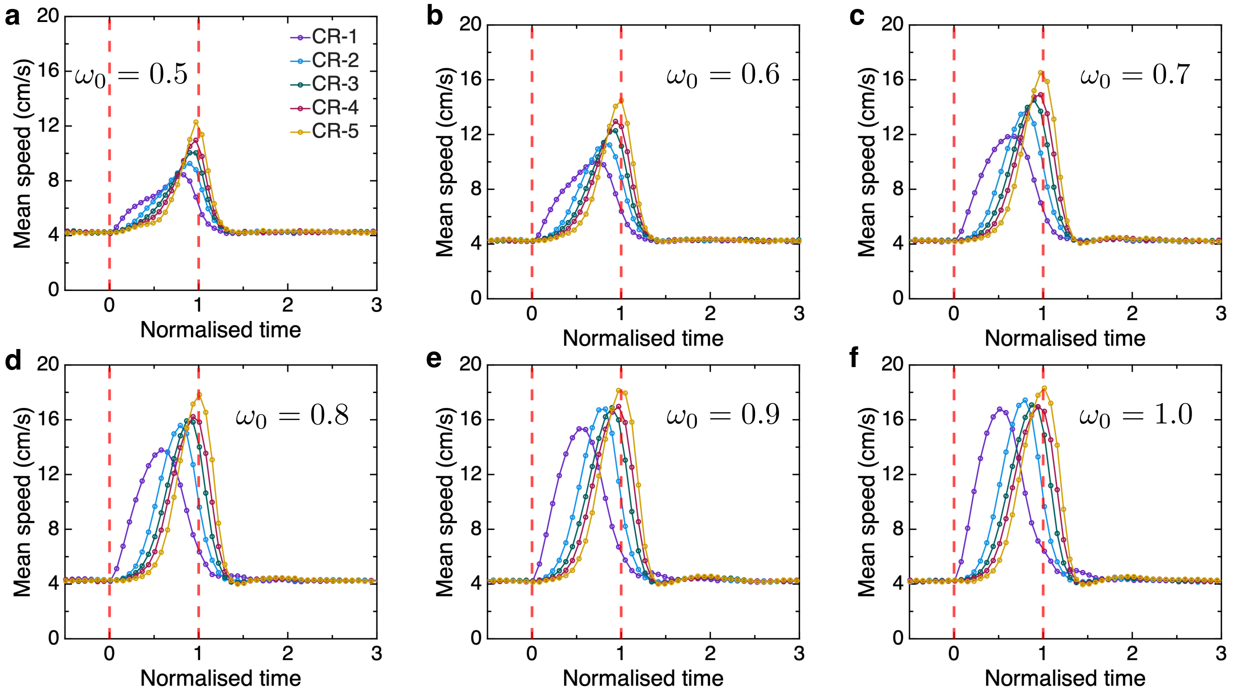


**Figure S10. Influence of escape-force strength on speed.** Mean swimming speed for simulations with (**a**) $\omega_0 = 0.5$, (**b**) $\omega_0 = 0.6$, (**c**) $\omega_0 = 0.7$, (**d**) $\omega_0 = 0.8$, (**e**) $\omega_0 = 0.9$, and (**f**) $\omega_0 = 1.0$. Increasing the initial escape-force strength alters the sequential speed dynamics associated with hurdle crossing.

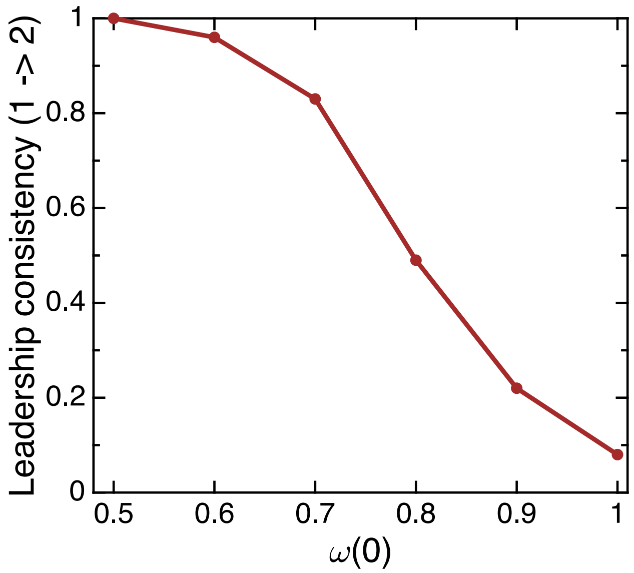


**Figure S11. Effect of $\omega(0)$ on the structure of the leadership network.** Fraction of statistically significant simulations in which the fish with crossing rank 2 follows the conditioned fish, plotted as a function of $\omega(0)$.

### Supplementary Tables

| Fish crossing rank | Possible edges | Observed edges in data | Observed edges in model |
|---|---|---|---|
| 1 (Conditioned fish) | 1 → 2<br>1 → 3<br>1 → 4<br>1 → 5 | None | None |
| 2 | 2 → 1<br>2 → 3<br>2 → 4<br>2 → 5 | 2 → 1 | 2 → 1 |
| 3 | 3 → 1<br>3 → 2<br>3 → 4<br>3 → 5 | 3 → 2<br>3 → 4 | 3 → 2 |
| 4 | 4 → 1<br>4 → 2<br>4 → 3<br>4 → 5 | 4 → 2<br>4 → 3 | 4 → 2<br>4 → 3 |
| 5 | 5 → 1<br>5 → 2<br>5 → 3<br>5 → 4 | 5 → 3<br>5 → 4 | 5 → 2<br>5 → 3<br>5 → 4 |

**Table S1. Hierarchical leader–follower relationships across crossing ranks.** Summary of all possible and experimentally observed leader-follower pairs for the experimental data and the computational model (here, arrows point from leaders to followers). The resulting interaction structure reveals a hierarchical organisation consistent with the order of crossing ranks. In the computational model, only the leader-follower pairs exhibiting statistically significant correlations in at least 50% of simulations are shown. A detailed representation of the corresponding interaction network is provided in figure 4c of the main text.

| **Parameter** | **Description** | **Value** |
|---|---|---|
| Preferred distance to neighbours | $r_{\rm D}$ | $5\,\mathrm{cm}$ |
| Preferred distance to wall | $l_{\rm w}$ | $3\,\mathrm{cm}$ |
| Preferred speed | $\mathrm{v}_0$ | $4.8\,\mathrm{cm\,s^{-1}}$ |
| Speed relaxation | $\beta$ | $1.2\,\mathrm{s^{-1}}$ |
| Turn friction | $\alpha$ | $4\,\mathrm{cm\,s^{-1}}$ |
| Angular noise | $D_\phi$ | $19.2\,\mathrm{cm\,s^{-2}}$ |
| Velocity noise | $D_v$ | $19.2\,\mathrm{cm\,s^{-2}}$ |
| Alignment strength | $\mu_{\rm Ali}$ | $0.56\,\mathrm{s^{-1}}$ |
| Distance regulation strength | $\mu_{\rm Att}$ | $36.8\,\mathrm{cm\,s^{-2}}$ |
| Wall alignment strength | $\mu_{\rm w}$ | $480\,\mathrm{cm\,s^{-2}}$ |
| Escape strength | $\mu_{\rm esc}$ | $20\,\mathrm{cm\,s^{-2}}$ |
| Distance slope | $m_d$ | $2.5\,\mathrm{cm^{-1}}$ |
| Relative strength of escape force at $t = 0$ | $\omega_0$ | $[0.5, 0.6, 0.7, 0.8, 0.9, 1]$ |
| Decay rate of escape force | $\gamma$ | $1\,\mathrm{s^{-1}}$ |

**Table S2. Parameters used in the computational escape model.** Parameter values were selected such that simulated individuals exhibited swimming speeds comparable to those observed experimentally during the initial and relaxation phases.
